# *sierra-local*: A lightweight standalone application for secure HIV-1 drug resistance prediction

**DOI:** 10.1101/393207

**Authors:** Jasper C Ho, Garway T Ng, Mathias Renaud, Art FY Poon

## Abstract

Genotypic resistance interpretation systems for the prediction and interpretation of HIV-1 antiretroviral resistance are an important part of the clinical management of HIV-1 infection. Current interpretation systems are generally hosted on remote webservers that enable clinical laboratories to generate resistance predictions easily and quickly from patient HIV-1 sequences encoding the primary targets of modern antiretroviral therapy. However they also potentially compromise a health provider’s ethical, professional, and legal obligations to data security, patient information confidentiality, and data provenance. Furthermore, reliance on web-based algorithms makes the clinical management of HIV-1 dependent on a network connection. Here, we describe the development and validation of *sierra-local*, an open-source implementation of the Stanford HIVdb genotypic resistance interpretation system for local execution, which aims to resolve the ethical, legal, and infrastructure issues associated with remote computing. This package reproduces the HIV-1 resistance scoring by the web-based Stanford HIVdb algorithm with a high degree of concordance (99.997%) and a higher level of performance than current methods of accessing HIVdb programmatically.

## INTRODUCTION

Genotype-based prediction of human immunodeficiency virus type 1 (HIV-1) drug resistance is an important component for the routine clinical management of HIV-1 infection [1, 2]. Detecting the presence of viruses carrying mutations that confer drug resistance enables physicians to select an optimal drug combination for that patient’s treatment regimen. Furthermore, genotyping by bulk sequencing is a cost-effective alternative to the direct measurement of drug resistance from culturing virus isolates in a laboratory [3]. Provided access to affordable bulk sequencing at an accredited laboratory for clinical microbiology, the interpretation of HIV-1 sequence variation is the primary obstacle to utilizing resistance genotyping for HIV-1 care. Fortunately, there are several HIV-1 drug resistance interpretation algorithms that can be accessed at no cost through web applications or services hosted by remote network servers, such as the Standard University HIV Drug Resistance Database (HIVdb) [4], Agence Nationale de Recherche sur le SIDA (ANRS) AC11 [5], and Rega Institute [6] algorithms. The Stanford HIVdb interpretation system can be accessed either through a web browser at http://hivdb.stanford.edu/hivdb or programmatically through its Sierra Web Service [7], which requires the transmission of an HIV-1 sequence from a local computer over the network to the remote server. This is a convenient arrangement for clinical laboratories because there is no need to install any specialized software, web browsers are ubiquitous and most users are familiar with submitting web forms.

On the other hand, there are a number of disadvantages to accessing interpretation systems over a network connection. First, HIV-1 sequences are sensitive patient information, not only because infection with HIV-1 remains a highly stigmatized condition, but also because sequence data have been used as evidence in the criminal prosecution of individuals for engaging in sexual intercourse without disclosing their infection status, leading to virus transmission [8]. Once sequence data have been transmitted to a remote server, one cedes all control over data security. Preventing the onward distribution of the data and deleting the data once the analysis is complete, for instance, is entirely the responsibility of the system administrators of the host server. Furthermore, unless the host server employs a secure transfer protocol, the unencrypted data are transmitted in the clear between a number of intermediary web servers, exposing these data to a ‘man-in-the-middle’ attack [9].

Second, the algorithm hosted on the server is effectively a black box — one has no insight into how resistance predictions are generated. Even if a version of the algorithm has been released into the public domain, one cannot be certain that the exact same algorithm was applied to their transmitted data. Importantly, different versions of a given algorithm can output significantly different resistance predictions, with the general trend being an increase in both resistance scores and predicted resistance levels [10]. In addition to contributing to inconsistencies in algorithm outputs, this makes it difficult to track data provenance, *i.e.*, the historical record of data processing, that has become recognized as a critical gap in the workflows of clinical laboratories. For instance, the College of American Pathologists recently issued new accreditation requirements stipulating that clinical laboratories must track the specific version of software programs used to process patient data [11]. Thus, a reliance on web-based systems creates significant issues for the reproducibility and quality assurance of clinical workflows. The Stanford HIVdb web service (Sierra [7]), for instance, automatically utilizes the most recent version of the HIVdb algorithm. While this constraint ensures that users employ the most up-to-date algorithm, it also introduces hidden changes to clinical pipelines, which may have been locally validated on older versions of the algorithm.

Third, dependence on a web resource may cause problems when the laboratory cannot access the host server, either due to local or regional network outages, or because the host server is malfunctioning or offline. In our experience, the web servers hosting the more popular HIV drug resistance interpretation algorithms such as the Stanford HIVdb database are reliable and well-maintained. However, it is not unusual for other web-based algorithms to be relocated or go offline when the developers move to other institutions or lack the resources to maintain the service.

One of the important features of the Stanford HIVdb algorithm is that it is regularly updated and released into the public domain in a standardized XML-based interchange format — the Algorithm Specification Interface version 2 (ASI2) format [12] — that was formulated and published by the same developers in conjunction with the Frontier Science Foundation. Here, we describe the implementation and validation of *sierra-local*, an open-source Python package for local execution of the HIVdb algorithm in the ASI2 format. This package utilizes, but does not require, a network connection to synchronize its local ASI2 file and reference data with the latest releases on the Stanford HIVdb web server. Our objective was to release a lightweight alternative to transmitting HIV-1 sequences to the HIVdb web server that minimizes the number of software dependencies, and that produces the exact same interpretations as the Sierra web service for all available HIV-1 sequences in the Stanford database.

## MATERIALS AND METHODS

Data Collection. We obtained the entirety of the genotype-treatment correlation datasets available through the Stanford HIV Drug Resistance Database (HIVdb [13]) on May 7 2018. These data included nucleotide sequences of 105,694 protease (PR) isolates, 112,723 reverse-transcriptase (RT) isolates, and 12,332 integrase (IN) isolates for a total of 230,749 sequences. In addition to sequence data, each record comprised of a list of the specific antiretrovirals (ARVs) that each isolate had been exposed to *in vivo* prior to collection, the region and year of collection, and subtype as determined by the Stanford University HIV Drug Resistance Database’s HIV Subtyping Program. After screening for empty and invalid data, the resulting dataset contained 103,711 PR entries, 110,222 RT entries, and 11,769 IN entries totalling 226,702 records. In addition, we retrieved 7 population-based HIV-1 *pol* datasets from Genbank using the NCBI PopSet interface (http://www.ncbi.nlm.nih.gov/popset). These datasets were selected from the most recent uploads of substantial numbers of HIV-1 sequences covering the regions encoding both PR and RT, and representing a diversity of HIV-1 subtypes and sampling locations around the world.

### Local HIVdb Algorithm Implementation

The team at Stanford HIVdb have created a Python tool, SierraPy (https://github.com/hivdb/sierra-client/tree/master/python), that serves as a command-line interface (CLI) for HIVdb. SierraPy does not process sequences directly, however, and only serves as a front-end for the HIVdb Sierra GraphQL Web Service (https://hivdb.stanford.edu/graphql) [7]. Its reliance on an active network connection to offload sequence processing to a remote server does not fulfill the usage gap we aim to address with *sierra-local*. To fill this gap, a system that provides a complete interface to a local version of the algorithm is needed. This local algorithm is first obtained as a publicly available HIVdb ASI2 file, which encodes both the algorithm for resistance scoring sequences and annotations describing relevant drug resistance mutations (DRMs) and ARVs of interest [12]. In short, it serves as a container for the core of the HIVdb resistance interpretation system which is not directly usable in a data-processing pipeline, as it is essentially only a descriptive XML defining the rules by which sequences should be scored. In the existing HIVdb pipeline, a Java-based interpreter generator called SableCC is used to compile an algorithm interpreter from the HIVdb ASI2 file, but we have not been able to find any such compiler in Python. The usage of SableCC in our local implementation of the Stanford algorithm would introduce further dependencies and obfuscate the clear relationship between the algorithm file and how local interpretations are generated. Hence, in light of the need for a Python-based local interpreter for the ASI2 format, we developed a regular expressions-based keyword-parsing method by which *sierra-local* locally compiles an executable model in Python directly from the local algorithm file. This method iterates through the HIVdb algorithm as an XML tree object in Python 3 and extracts the information encoded within using ASI2 keywords defined by the ASI2 Document Type Definition (DTD). *sierra-local* then uses this method to calibrate the model by assigning the drug clause definitions, drug class lists, resistance level interpretations, DRM comments, and complex drug-DRM scoring conditions to a set of dictionary and list objects. Once populated, this model serves as the framework for sequence resistance scoring.

### Sequence Pre-Processing and Validation

Prior to scoring, the HIVdb Sierra Web Service performs several pre-processing and validation steps on submitted query sequences and identified mutation sites found within sequences. We emulated these steps to maximize fidelity to the HIVdb pipeline, including sequence alignment, gene identification, mutation site classification, sequence trimming, sequence subtyping, and sequence validation. These pre-processing steps can be considered parts of the algorithm involved in generating resistance prediction scores that are not included and distributed in the HIVdb ASI2 file.

#### Sequence Alignment

Of particular importance in these steps is sequence alignment. HIVdb utilizes NucAmino [14] to initially align and identify amino acid mutations in each query sequence. This nucleotide-to-amino acid alignment program is optimized for viral gene sequences, and was developed by the HIVdb developers in the Go language. However, in practice, NucAmino does not return the aligned sequences themselves; instead it only returns a list of mutations relative to the consensus subtype B amino acid sequence and general sequence metadata, *e.g.*, the aligned start and end coordinates of the query sequence relative to HIV-1 *pol*. If *sierra-local* does not align the query viral sequences exactly as how HIVdb would, the mutations identified relative to the consensus subtype B sequence would not be identical in all cases. Since these mutations serve as the basis for resistance predictions, an identical alignment process is of the utmost importance in maintaining fidelity to HIVdb Sierra. Thus, we decided to incorporate the NucAmino alignment program as a dependency, rather than substituting a more integrated native implementation in Python. *sierra-local* calls a pre-compiled NucAmino binary as a Python subprocess with default settings to execute this pre-processing step. We used NucAmino version 0.1.3 for our validation experiments. The optional JSON output from NucAmino was captured in Python by redirecting the standard output stream from the subprocess.

#### Gene Identification

A critical part of resistance scoring is knowing which gene products encoded by HIV-1 *pol* (protease, RT and integrase) are present in the sequence being analyzed. To map the query sequence to these targets, we compared the aligned start and end positions returned by NucAmino to the HXB2 reference positions. For consistency with the HIVdb pipeline, amino acids were renumbered relative to the start position of the corresponding gene product (PR, RT and IN).

#### Mutation Site Classification

In the process of mutation site classification, each amino acid mutation site identified by NucAmino was further categorized as an insertion, deletion, or mixture using the original nucleotide sequence as a reference for amino acid translation. For consistency, we ported a Java method from the Sierra algorithm for determining nucleotide codon translation ambiguity to Python. Each site of interest identified by NucAmino on the aligned and translated query sequence generated a list of characters representing possible encoded amino acids at that site. Associating mutation sites with more than one encoded amino acid allows for a ‘fuzzy’ matching of a single sequence to multiple scoring conditions sharing the same residue position but different mutations. In certain cases, ‘highly ambiguous’ sites encoding more than four possible amino acids – made possible due to the presence of ambiguous nucleotides such as ‘R’ (A/G), ‘B’ (C/G/T), and ‘N’ (A/T/C/G) – were flagged as ‘ambiguous’ and represented with a translation of ‘X’.

#### Sequence Trimming

Leading and trailing regions in each sequence containing a minimum proportion (30%) of ‘low quality’ sites — defined as sequenced sites that are highly ambiguous, stop codons, unusual mutations, or frameshifts — with at least one of these sites every 15 residues were trimmed prior to resistance scoring. Based on our inspection of the Sierra source code, we defined a site as sequenced if the codon does not have more than one unknown nucleotide. Sites further qualified as ‘unusual mutations’ if they were indicative of APOBEC-mediated G-to-A hypermutation [15] or if the highest frequency of that mutation in the pooled untreated and treated viruses for that specific group M subtype was less than 0.1% in the Stanford University HIV Drug Resistance Database. We configured the *sierra-local* installation process to obtain a local copy of the reference data for APOBEC-mediated G-to-A hypermutations in HIV-1 *pol* and for other HIV-1 mutation prevalences from the Stanford HIVdb server, and to automatically update this local copy to accommodate changes in these reference data over time.

#### Sequence Subtyping

The previously discussed trimming step requires that sequences be subtyped in order to determine the frequency of mutations in subtype-specific pooled untreated and treated viruses. We wrote a Python implementation of HIVdb’s HIV Subtyping Program, which categorizes submitted sequences as a pure subtype, a circulating recombinant form (CRF), a unique recombinant form (URF), non-group M HIV-1, or HIV-2. This process calculates uncorrected pairwise distances (the proportion of nucleotide differences) between submitted nucleotide sequences and a set of 200 different subtype-specific reference sequences. Because the HIVdb HIV Subtyping Program is very simple and does not perform phylogenetic analysis or bootstrapping, its results may not be as accurate as more sophisticated systems more commonly used such as the Rega Institute HIV-1 Automated Subtyping Tool [4]. Comparing interpretation systems, it has been suggested that ANRS AC11, HIVdb, and Rega demonstrate discordances that may be subtype-dependent [16, 17]. Yet, current genotypic resistance interpretation systems, including Stanford HIVdb, are subtype-agnostic within themselves, meaning that they do not offer differential resistance penalty scoring based on the subtype identified. Thus, other than being used to trim submitted sequences, the subtyping in this system does not significantly influence HIV-1 sequence interpretation.

### Resistance Scoring

The HIVdb algorithm begins the scoring process by assessing each sequence’s potential resistance to 22 commonly-used ARVs independently by searching for ARV-specific mutation conditions present in the query sequence. ARV-specific mutation conditions encode and define the circumstances in which a mutation in a particular gene has been shown to influence resistance, and quantifies this resistance with an integer penalty score. Each of these mutation conditions are associated with one or more specified amino acid changes at one or more particular positions, the ARV of interest, and the gene of interest in HIV-1 *pol* targeted by the ARV. For a mutation condition’s resistance score to be counted, all of these criteria must be fulfilled.

*sierra-local* iterates over the ARV-specific mutation conditions corresponding to the gene regions detected in the query sequence. HIVdb Sierra also validates sequences and returns a list of validation problems found in each sequence. A sequence is invalidated or flagged as a query if: no genes are found; the sequence is a reverse complement; the genes have not been aligned properly; it is too short based on gene-specific minimum cutoff nucleotide lengths; it is trimmed at the 5’ or 3’ ends due to low-quality leading or trailing sites; an indel is longer than 30 base pairs; invalid ‘NA’ characters are found; one or more stop codons are found; too many ‘unusual mutations’ as previously described are found; the virus is subtyped as HIV-2; the number of APOBEC mutations is two or greater; the number of APOBEC mutations at drug resistance positions (DRPs) is one or greater; or if the count of frameshifts, unusual insertions, and unusual deletions together is positive. We emulated these validation steps in our pipeline to maximize parity with Sierra.

### Algorithm Output Generation

Once all queries are pre-processed, scored, and otherwise annotated, *sierra-local* writes these results into a JSON (JavaScript Object Notation) format that mimics the standard output format of the HIVdb Sierra Web Service. For the sake of brevity and simplicity, we decided to have *sierra-local* omit the ‘pretty pairwise’ sequence output found in Sierra’s standard output. This output format usually contains a numerical sequence of all residue positions, a reference amino acid sequence, an aligned nucleotide sequence, and a mutation-only list. These data are not critical to a rapid genotypic interpretation system and may be omitted without detriment to the central aim. Furthermore, their exclusion from the results leads to a five-fold reduction in the results file size, which may be significant when processing large batches of sequences.

### Validation Against HIVdb Sierra

We scored the entirety of the partitioned and filtered HIVdb genotype-treatment correlation dataset as of May 7 2018 with both *sierra-local* and SierraPy (version 0.2.1), storing all output results files from both programs. Validation was conducted using the HIVdb version 8.5 algorithm on both platforms. Because the algorithm was updated to version 8.6.1 during the validation experiments, we used the newer version for the HIV-1 integrase data sets since the update mostly affected the interpretation of mutations within this region. Subsequent analysis of validation testing was conducted in R (version 3.4.4) [18] using the *jsonlite* package (version 1.5) [19] to extract resistance scores and relevant meta-data from the JSON results files from either program. For each sequence record, the resistance scores from the pipelines were compared for identicality. Validation and mutational metadata for discordant cases were analyzed for iterative refinement of the *sierra-local* scripts until a satisfactory level of concordance was attained. Subsequently, we carried out a second set of validation experiments on longer HIV-1 *pol* (PR-RT) sequence data sets under HIVdb algorithm version 8.7.

### Performance Measurements

To quantify the speed of *sierra-local* in processing sequences, we chose a random subsample of gene-specific batches balanced across the genes of HIV-1 *pol* in our filtered genotype-treatment correlation dataset. This sampled dataset comprised of 9,673 PR sequences, 10,000 RT sequences, and 9,720 IN sequences contained in 10 batches per gene. Each gene-specific batch, roughly 10,000 sequences in size, was independently processed with both *sierra-local* and SierraPy. This method was timed using the *time* package in Python to determine file run-times and the resulting processing speeds of each program, measured in sequences per second.

### Software Availability

The source code for *sierra-local* has been released under the GNU General Public License (version 3) and may be obtained at http://www.github.com/PoonLab/sierra-local or from the Python Package Index (https://pypi.org/project/sierralocal). Detailed installation instructions are provided on the GitHub website.

## RESULTS AND DISCUSSION

### Implementation

Distribution of the Stanford HIVdb algorithm in an XML-based interchange format called the Algorithm Specification Interface version 2 (ASI2) enables a seamless approach to updating the algorithm and algorithm version control. Despite the accessibility of the HIVdb algorithm itself, the ASI2 file is necessary but not sufficient to generate resistance predictions. Additional steps needed but not encoded by the algorithm file include: sequence alignment, sequence quality control and validation, sequence trimming, sequence subtyping, and formatting of the results output file with resistance predictions and accessory meta-data. These processes comprise the bulk of the functionality developed in *sierra-local* and are coordinated to generate resistance predictions in a manner as identical as possible to HIVdb Sierra.

As new versions of HIVdb are released to reflect the growing knowledge of HIV resistance, users may easily manually update their local copy of the algorithm with a provided Python script (updater.py). We also provide the option to automatically update the algorithm to the most recently released version at run-time. For example, three major updates to the HIVdb algorithm were released to the public during the development of *sierra-local*. Our local copies of these files were automatically retrieved by the *sierra-local* pipeline, but we also configured the pipeline to use specific versions by setting the ‘-xml’ option. The option to choose between automatic updating or freezing the algorithm to a specific version enables physicians and researchers to fulfill potential version tracking and data provenance requirements. However, even with functionalities addressing these obligations embedded in *sierra-local*, the health care provider still bears the responsibility of operating with a knowledge of their software and algorithm versions and the changes between these.

### Introduction

Concordance with HIVdb Sierra. Out of the 103,711 PR, the 111,222 RT, and the 11,769 IN sequences (total 226,702) processed with both *sierra-local* and HIVdb SierraPy pipelines, the predicted resistance scores and component subscores were completely identical in 226,696 (99.997%). Of the 6 sequences that did not have identical scores for all ARVs between the pipelines, 3 were PR sequences and 3 were RT (Table 1). The most frequent cause of discordance was the trimming of nucleotides in the leading or trailing ends of the sequence on the basis of the prevalence of the amino acid polymorphism in the corresponding HIV-1 subtype or ambiguous base calls.

**Table 1.** Discordant cases between *SierraPy* and *sierra-local*. Both pipelines were applied to the same database, comprising 103,711 PR, 110,222 RT and 11,769 IN records. We observed a total of 3 discordant cases in PR, 3 cases in RT, and none in IN. For cases where both pipelines generated resistance score predictions, we listed the discordant scores for the respective drug names. Putative reasons for discordance were assessed from validation outputs.

| Accession | Gene | Length (nt) | Mixtures | Reason | Drug | *SierraPy* | *sierra-local* |
| --- | --- | --- | --- | --- | --- | --- | --- |
| AY739171 | PR | 279 | 15 | No genes found |  |  |  |
| GU188744 | PR | 216 | 0 | Sequence trimming, unusual mutations | ATV/r | 0 | 5 |
|  |  |  |  |  | DRV/r | 0 | 5 |
|  |  |  |  |  | FPV/r | 0 | 10 |
|  |  |  |  |  | IDV/r | 0 | 5 |
|  |  |  |  |  | LPV/r | 0 | 5 |
|  |  |  |  |  | NFV | 0 | 10 |
|  |  |  |  |  | SQV/r | 0 | 5 |
|  |  |  |  |  | TPV/r | 0 | 10 |
| JQ028402 | PR | 243 | 0 | Sequence trimming | ATV/r | 0 | 5 |
|  |  |  |  |  | FPV/r | 0 | 5 |
|  |  |  |  |  | IDV/r | 0 | 5 |
|  |  |  |  |  | NFV | 0 | 15 |
|  |  |  |  |  | SQV/r | 0 | 5 |
| KT745612 | RT | 630 | 10 | Stop codon | ABC | 100 | 95 |
|  |  |  |  |  | AZT | 130 | 125 |
|  |  |  |  |  | d4T | 130 | 125 |
|  |  |  |  |  | DDI | 100 | 95 |
|  |  |  |  |  | TDF | 70 | 65 |
| DQ297313 | RT | 564 | 0 | Sequence trimming | ABC | 0 | 5 |
|  |  |  |  |  | AZT | 0 | 20 |
|  |  |  |  |  | d4T | 0 | 20 |
|  |  |  |  |  | DDI | 0 | 10 |
|  |  |  |  |  | TDF | 0 | 5 |
| KC221011 | RT | 597 | 14 | Sequence trimming | ABC | 10 | 25 |
|  |  |  |  |  | AZT | 25 | 50 |
|  |  |  |  |  | d4T | 35 | 60 |
|  |  |  |  |  | DDI | 45 | 65 |
|  |  |  |  |  | TDF | 10 | 25 |

We examined the distributions of sequence lengths and mixtures (ambiguous base calls) in the database to determine whether the discordant cases might be explained by these factors. Of the three discordant PR sequences, AY739171 contained an extremely large number of mixtures (15). In addition, this sequence was derived from an HIV-1 group O infection. Only 0.42% of PR sequences in the databases contained as many or more mixtures; the median [2.5% and 97.5% quantiles] number of mixtures was 1 [0, 9]. The remaining two PR sequences were unusually short, where the median length was 297 nt; the proportion of sequences with lengths shorter than GU188744 and JQ028402 were 0.43% and 0.76%, respectively. Although *sierra-local* reported non-zero resistance scores where *SierraPy* reported none, the scores were generally in the range of susceptible to potential low-level resistance interpretations. Similarly, the three discordant RT sequences were either short (overall median [quantiles] = 774 [588,1680] nt) or contained substantial numbers of mixtures for sequences of comparable length (*e.g.*, 4 [0, 19] mixtures for sequences between 550 and 650 nt in length). In the latter case, however, neither KT745612 nor KC221011 contained a significantly excessive number of mixtures. These three RT sequences also resulted in slighty more discordant resistance interpretations; for example, the d4T resistance score for KC221011 was switched by *sierra-local* from intermediate to high-level resistance.

In addition, we ran both pipelines on six recently published sets of HIV-1 *pol* sequences comprising both PR and RT encoding regions. These data sets were selected to cover a diversity of HIV-1 subtypes and locations around the world (Table 2). The major HIV-1 group M subtypes A, B, C and D were represented in these data, as well as several circulating recombinant forms (CRFs) such as CRF07_BC, which is highly prevalent in East Asia. All resistance scores for all 1,006 sequences were completely concordant between the pipelines.

**Table 2.** Characteristics of HIV PR-RT population data sets. Sequence data were obtained for an arbitrary selection of recent studies with HIV-1 *pol* sequences deposited in Genbank that spanned both PR and RT. We processed the sequences through both *SierraPy* and *sierra-local* to confirm that the pipelines obtained identical resistance scores. Subtypes listed were obtained from the *SierraPy* pipeline; the subtype classifications obtained from *sierra-local* were highly concordant (94.5% identical). CRF = circulating recombinant form.

| Country/region | Sample size | Subtypes | Sequence length (nt) | Accession numbers | Ref. |
| --- | --- | --- | --- | --- | --- |
| Brazil | 103 | B (100%) | 1262.0 | MF545238 – MF545340 |  |
| Ethiopia | 67 | C (97.0%), B (1.5%), A (1.5%) | 1042.1 | MH324937 – MH325003 | [20] |
| Guinea-Bissau | 54 | CRF02_AG (88.9%), A (5.6%), CRF06_cpx (1.8%) | 1035.0 | MH605452 – MH605505 | [21] |
| Hong Kong | 284 | C (36.0%), CRF07_BC (36.0%), CRF02_AG (8.8%) | 1157.8 | MH757122 – MH757405 |  |
| South Africa | 212 | C (100%) | 1195.0 | MH920641 – MH920852 | [22] |
| Tajikistan | 146 | A (97.3%), CRF02_AG (2.0%), CRF63_02A1 (0.7%) | 1351.1 | MH543115 – MH543260 |  |
| Uganda | 140 | D (99.3%), CRF10_CD (0.7%) | 864.0 | MH925538 – MH925677 |  |

### Increased Performance over HIVdb Sierra

Performance and hence, the speed, of software packages varies according to hardware, software, and input data characteristics. All development, testing, and validation was performed on a workstation running Ubuntu 18.04 LTS with an Intel Xeon E5-1650 v4 hexa-core CPU at 3.60 GHz and 16 GB of DDR4-2400 RAM with a gigabit network connection. *sierra-local* achieved mean [range] processing speeds of 47.08 sequences/second (seq/s) [45.07, 48.49] for PR, 16.20 seq/s [14.01, 19.97] for RT, and 14.99 seq/s [14.79, 15.56] for IN. A substantial fraction of processing time was consumed by subtyping. SierraPy, with the same dataset as previously described, yielded mean processing speeds of 16.01 seq/s [12.88, 17.60] for PR, 6.12 seq/s [4.83, 7.54] for RT, and 5.19 seq/s [5.05, 5.47] for IN. Although the size of sequence batches used in this performance comparison likely is a factor in the results by virtue of file writing and reading being done once per batch, the large batch size used minimizes the effect of these I/O processes on the overall runtime. Overall, *sierra-local* is able to process and return results for submitted query HIV-1 *pol* sequences roughly 3 times faster than SierraPy, depending on the nature of the sequences and the type of local computing resources available. This result is in the expected direction since local computing resources are able to be fully utilized, whereas SierraPy depends on server-side processing speed, server load, request balancing, as well as network speed and traffic. In this case, network speeds between SierraPy on the local workstation and HIVdb Sierra GraphQL Web Service were likely not a significant factor in the speed improvement results obtained. With slower network speeds and all other factors being equal, however, the relative processing speed of *sierra-local* to SierraPy can only be expected to increase.

### Concluding remarks

The distribution of the HIVdb resistance genotyping algorithm in a standardized format (ASI [12] is an important resource for HIV-1 research and clinical management, and an exemplary case of open science. *sierra-local* provides a convenient framework to generate HIV drug resistance predictions from ASI releases in a secure environment and confers full control over data provenance. The ability to apply ASI-encoded algorithms locally (offline) also makes this part of the laboratory workflow robust to network availabilty may be particularly important for laboratories situated in resource-limited settings. In addition, the relative processing speed of *sierra-local* can confer an advantage for research applications requiring the analysis of large numbers of sequences. The emulation of an established genotype interpretation system to process unaligned nucleotide sequences and produce identical resistance predictions and data summaries in a small, standalone package was not a trivial undertaking. Despite the relative simplificity of the rules-based HIVdb algorithm, there were a large number of pre-processing and post-processing steps that were necessary to adapt to maximize concordance with the original system. We developed *sierra-local* with the aims of minimizing the number of additional programs that users would have to install for a local implementation. The only other presently available means for implementing for a completely independent instance of the HIVdb algorithm is by hosting an instance of the HIVdb Sierra Web Service itself, which was recently made possible with the release of the Sierra source code. This approach, however, requires the configuration of a web server, the Apache Tomcat web container, and a large number of Java libraries (Apache Commons Lang, Apache Commons Math, Apache Commons IO, Apache Log4j, Google Guava, Google Gson, protonpack, and GraphQL-Java). Furthermore, hosting a stable web service for the sole purpose of independently generating clinical resistance predictions increases facility requirements for servers, information technology support staff, and generally complicates an already complex workflow. We hope that making this lightweight, open-source implementation of the HIVdb ASI to the clinical and research community will further democratize HIV drug resistance genotyping across providers of HIV care.

## ACKNOWLEDGEMENTS

We thank Philip Tzou for bringing NucAmino to our attention, and for his contributions to open science in the release of the Stanford HIVdb resistance program source code. This work was supported in part by the Government of Canada through Genome Canada and the Ontario Genomics Institute (OGI-131) and by a grant from the Canadian Institutes of Health Research (PJT-156178). The funders had no role in study design, data collection and interpretation, or the decision to submit the work for publication.

